# Reassessing the Neural Correlates of Social Exclusion: A Replication Study of the Cyberball Paradigm Using Arterial Spin Labelling

**DOI:** 10.1101/2024.08.08.607136

**Authors:** Karin Labek, Roberto Viviani

## Abstract

**Background/Objectives:** The cyberball paradigm has been used in numerous neuroimaging studies to elicit activation in neural substrates of social exclusion, which have been interpreted in terms of activity associated with “social pain”. The objectives of the study were to assess not only the replicability but also the specificity of the areas activated by this paradigm.

**Methods:** Functional imaging with arterial spin labelling, an approach to image longer mental states.

**Results:** we replicated findings of previous meta-analyses of this paradigm in the inferior frontal gyrus and ventral cingular cortex. However, these areas were also active in a watch condition (in which participants were not excluded), although less so.

**Conclusions:** These findings relativize a simple and specific interpretation of these areas as the neural substrates of social exclusion and social pain, as in previous studies. In a broader experimental context, similar activations have been reported by neuroimaging studies when semantic disambiguation and evaluation of action goals are required, an interpretation that may apply also to the effects elicited by this paradigm.

## Introduction

Social pain has been described as a distressing emotional experience resulting from rejection, loss, or social exclusion. Everyone has experienced it, making its understanding not solely a scientific question but also a universal human concern, which qualifies it as a transdiagnostic phenomenon. Several paradigms have been developed to study social pain using fMRI, each providing unique insights into its neuronal underpinnings in healthy and patients suffering from mental health problems. Examples include trust games (King-Casas et al. 2008), social feedback paradigms (Somerville et al. 2006; Masten et al. 2011), rejection simulations (Kross et al. 2011; Eisenberger et al. 2003), and passive exposure to loss (Labek et al. 2017).

Among these, the cyberball paradigm (Williams et al. 2000) has been frequently investigated in neuroimaging studies. It is a virtual ball-tossing game where participants experience stages of inclusion and exclusion. In a first practicing phase, participants view two virtual players on a computer screen exchanging a ball (watch condition). In a second phase, the ball is exchanged with the participant (play condition). In a third and final phase, the participant is unexpectedly excluded from play from the players (exclusion condition). This paradigm became notorious after Eisenberger et al. (2003) showed in a functional imaging study the activation of the dorsal anterior cingulate (dACC, an area associated with pain in previous studies, Davis et al. 1997; Sawamoto et al. 2000). This finding led to the proposal of the “social pain” hypothesis, which suggests that social and physical pain share common neural mechanisms/activations (Panksepp 2003; Eisenberger et al. 2003).

Subsequent studies carried out with this paradigm have considerably expanded the areas associated with exclusion, while often failing to replicate the original finding. Cacioppo et al. (2013) failed to find significant dACC effects, instead noting activations in the ventromedial prefrontal cortex/ventral anterior cingular cortex (vACC), anterior insula (aI), and lateral orbitofrontal cortex/inferior frontal gyru (OFC/iFG). Vijayakumar et al. (2017) replicated these results, adding to them the posterior cingulate cortex (pCC), with no significant dACC activation. They also reported activation within a left PFC cluster that includes the ventrolateral PFC and the lateral OFC, extending to the left IFG. A meta-analysis by Rotge et al. (2014) demonstrated engagement of both ventral and dorsal subdivisions of the ACC, with the subgenual and pregenual vACC particularly associated with self-reported distress during social exclusion (Rotge et al. 2014).

While the precise findings of these meta-analyses varied, they question the role of dACC for social pain, instead drawing attention to other common themes. One is the recruitment of the medial portion of the default mode network (DMN), which includes the vACC and pCC. In their recent meta-analysis, Mwilambwe-Tshilobo and Spreng (2021) found that social exclusion reliably engages the medial DMN, while not reliably activating the dACC. The authors warn against attributing a function specifically related to social exclusion to the areas identified with this paradigm. A second area that is consistently reported in these meta-analyses is the iFG.

These subsequent findings justify a reassessment of the cyberball paradigm in at least two respects. The first is looking at its neural underpinnings in terms of the large-scale organization of the cortex and a broader experimental context. The DMN, which is coextensive with semantic association areas (Binder et al. 2009), is the terminal point of increasingly abstract multimodal encodings of external and internal representations (Mesulam 1990), and differs from unimodal association areas in displaying extensive long-range connectivity (Bassett and Bullmore 2006; Sporns et al. 2007; Margulies et al. 2016). Given that social cognition and emotion appraisal involve schema recruitment, it may not be surprising that these semantic areas are frequently active in social cognition tasks as the terminal point of progressively more abstract stimulus encoding (Messina et al. 2016; Viviani et al. 2020; Labek et al. 2023) and appear in general reviews of these neuroimaging studies (Schurz et al. 2021). The iFG, which has been proposed as a point of integration between these ventral areas and the dorsal network activated by cognitive effort (Corbetta et al. 2008), is also consistently activated when semantic disambiguation is needed (Demb et al. 1995; Thompson-Schill et al. 1997; Binder and Desai 2011), and by social cognition tasks where violations of social expectations impose a reassessment of the interaction (Sanfey et al. 2003; Montague and Lohrenz 2007; King-Casas et al. 2008). These neural substrates will be the focus of the present study, using a region of interest approach to improve sensitivity.

The second is looking at effects that were not considered in the original study by Eisenberger et al. (2003). In that study, the neural substrates of social exclusion were identified by comparing the exclusion condition with the condition in which participants were actively playing the game. However, it may be argued that the initial practice condition may be an even more appropriate control condition to identify exclusion. This is because in both cases the activity of participants is the same (watching the game), in one case becoming aware of the exclusion. In contrast, the playing condition may involve recruitment of the resources required for active play. Since increased cognitive recruitment may depress the DMN, the question arises of the extent to which both practice and exclusion conditions share a common DMN recruitment. Therefore, broadening the scope of the contrasts considered in the analysis might provide information relevant to place the findings within the large-scale cortical organization we have just mentioned.

Finally, our study differs from most other cyberball studies in the use of arterial spin labelling (ASL) perfusion MRI. ASL provides absolute quantification of cerebral blood flow (CBF), offering a direct measure of perfusion in specific brain regions. This quantitative approach allows for comparisons across different static conditions and is especially appropriate to image lang-lasting emotional states, as it does not require, as classic EPI-based imaging, relatively quick alternations of experimental and control conditions. Instead, ASL allows planning experiments as homogenous block sessions in which participants were exposed to a homogeneous condition, as in PET designs. This is particularly appropriate here since the exclusion condition is a protracted experience, which can be compared to homogenous play and watch conditions. Despite these advantages, only a few studies have utilized ASL in conjunction with the cyberball paradigm (Kiefer et al. 2021).

## Results

We first verified that the ASL technique was implemented successfully by looking at the contrast play vs. (watch or exclude), as we expected processes recruited during active play to be identified by this contrast. As expected, dorsal cortical areas involved in attentional processing (frontal eye fields, intraparietal gyrus) were active at significant peak and cluster level in this contrast (Figure 1, *z*=+48, red-yellow; *p* < 0.001, corrected at cluster-level; for other corrections not reported in text, see Table A1 in the Appendix). In the other direction we see areas that were recruited by the watch or the exclusion conditions, or both simultaneously (*z*=-6, blue-green, all significant *p* < 0.001, cluster-level corrected, Table A1). One can see that, in the medial face (x=-5 in Figure 1), they corresponded to areas of the DMN: the vACC and the pCC. This latter extended towards motor planning areas in premotor cortex.

**Figure 1.**
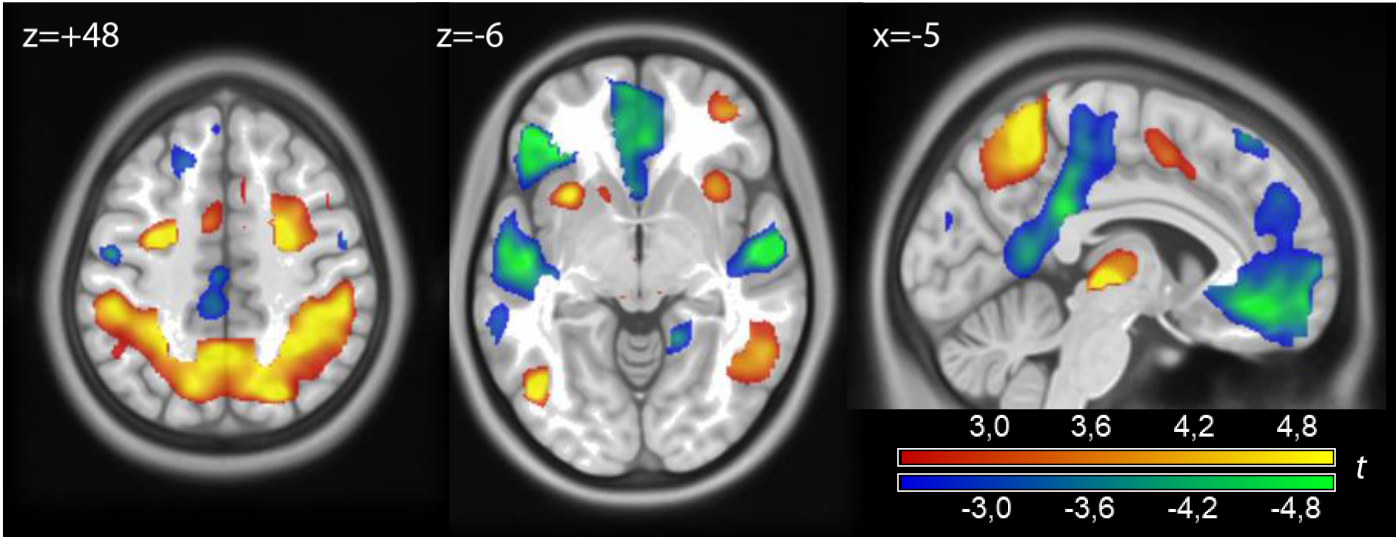
Contrast play vs. (watch or exclusion), in red-orange. In blue-green, effects in the opposite direction. Coordinates in Montreal Neurological Space. Parametric maps of *t* values displayed at *p* < 0.01, uncorrected.

We then looked at the exclusion vs. play contrast to see if we could replicate the findings of the cyberball paradigm in the literature (Figure 2, red colors, and Table A2 in the Appendix). Significant effects at the peak and cluster level were detected in the iFG (*p* = 0.002, cluster level-corrected) in a large cluster extending into the right-anterior portion of the middle temporal gyrus, and in the vACC (*p* = 0.018, cluster-level corrected; clusters #1 and #2 in Table A2). No significant effects were detected, even at uncorrected levels, in dACC. Activity in the pCC was present only at uncorrected levels and failed to reach significance.

**Figure 2.**
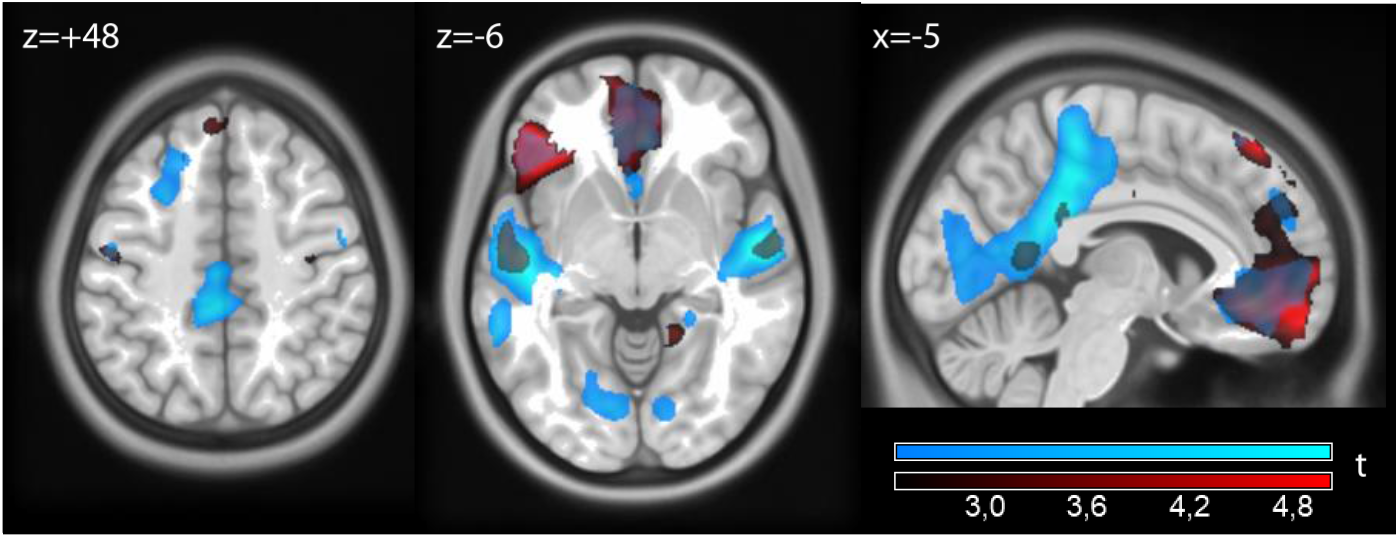
In red: contrast exclude vs. play. In light blue: contrast watch vs. play. Data displayed at *p* < 0.01, uncorrected.

We then looked at the contrast watch vs. play (Figure 2, light blue, and Table A3 in the Appendix). The largest effect here was in the anterior portion of the middle temporal gyrus bilaterally, on the left extending posteriorly toward Heschl’s gyrus (clusters #1 and #2 in Table A3, all *p* < 0.001, cluster-level corrected). One can also see an intense effect in the pCC, significant at peak and cluster level (cluster #3, *p* = 0.001), extending dorsally into premotor cortex. The vACC was also active at cluster level (cluster #4 in Table A3, *p* = 0.026, all cluster-level corrected).

One can see in Figure 2 that the effects of exclusion and watch overlapped. The main hubs of these effects (iFG, vACC, and anterior temporal lobe) were significant in both contrasts. It therefore appears that both contributed to the effects shown in green in Figure 1. However, there was a tendency for the exclusion contrast to involve preferentially prefrontal areas, whereas the watch contrast was most marked in posterior areas.

The last contrast we looked at was the contrast exclude vs. watch (Figure 3 and Table A4 in the Appendix). This contrast tested the significance of preferential distribution in anterior and posterior areas of the effects of exclusion and watch, relative to play. This contrast recorded the higher activity in the iFG and the anterior portion of the vACC of exclusion (clusters #2 and #3 in Table A4), although this effect was significant only at the more lenient corrections for these two regions of interest (*p* = 0.031 and *p* = 0.027, peak-level), in contrast to all other effects reported here. In the other direction (watch vs exclude), we found extensive effects in visual areas extending anteriorly towards the middle temporal gyrus (clusters #4 and #9 in Table A4). These areas were located posteriorly to the common effects of exclude and watch conditions in the temporal lobe and in the pCC, but adjacent to them.

**Figure 3.**
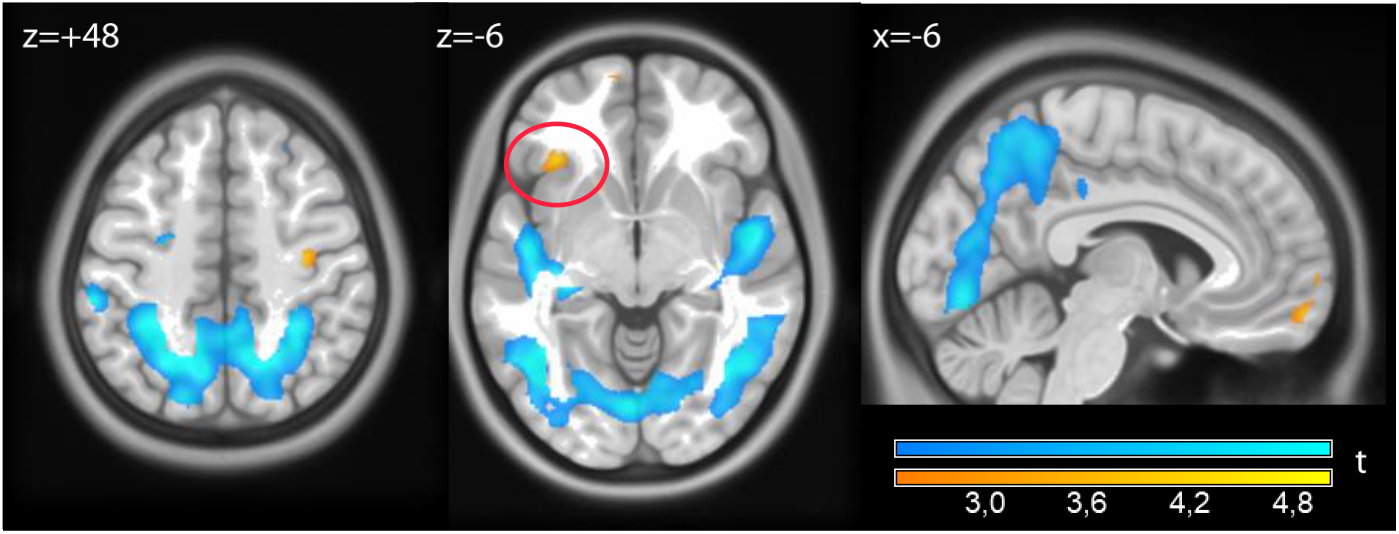
In yellows: contrast exclusion vs. watch. The red circle shows the left iFG/frontal operculum. In light blue: contrast watch vs. exclusion. Data displayed at *p* < 0.01, uncorrected.

## Discussion

Social pain refers to the distressing experience that results from social rejection, exclusion or loss. It encompasses the emotional pain resulting from social disconnection, a form of pain which, as it has been argued (Panksepp 2003), may have an evolutionary basis. However, Eisenberger’s study has faced significant criticism. Apart from the low replicability of the original dACC finding, critics have argued that activations could be related to general conflict detection or expectancy violation processes rather than social pain specifically (Cacioppo et al. 2013). They also suggest that these activations might be indicative of broader negative affective processes or the processing of salient events, rather than of a specific form of pain per se. A similar criticism has been formulated for the evidence for the shared activations mechanism (Decety 2010). This criticism highlights the difficulty of interpreting neuroimaging results in social exclusion paradigms as the neural correlates of specific socio-emotional processes and underscore the need for careful experimental design and interpretation in this field of research.

Our findings can be brought to bear on this criticism in two respects. First, the cortical modulations observed in the exclusion condition were relatively small perfusion changes affecting a network commonly modulated by both the watch and exclusion conditions, suggesting a shared functional role. This network was partially co-extensive to the DMN but did not include areas typically found as effects of task deactivations, such as the inferior parietal junction. In the posterior part of the brain, which is deputed to the visuospatial analysis of the environment, these areas were located far from primary and secondary visual areas (i.e., the anterior part of the middle temporal gyurs/temporal poles, and pCC). In previous studies, we have highlighted the role of homologous areas in emotion appraisal, showing that they are located at the terminal of a gradient of activity, associable to progressively abstract encodings, consistent with a function in schema recruitment (Viviani et al. 2020; Labek et al. 2023). One can see that in our data the temporal pole and pCC activity was located anteriorly to the visual encoding activity elicited by the watch condition, consistent with high-level encoding. This functional organization is generic, in the sense that it follows general principles of encoding of sensory information (Mesulam 1990). It has also been shown that these high-level association areas share long-range connectivity with the DMN, in contrast to lower-level unimodal association areas (Margulies et al. 2016; Viviani et al. 2020). It has therefore been suggested that the DMN constitutes a core cortical network with the capacity to relay information between the high association areas of the cortex (Bassett and Bullmore 2006; Sporns et al. 2007; van den Heuvel and Sporns 2013). The relative specialization of the recruited areas in the watch (pCC) and in the exclusion conditions (vACC) are consistent with the prevalent role of visual information in the former, and of information about action goals (Vogt et al. 1992; Arana et al. 2003; FitzGerald et al. 2012; Rudebeck and Murray 2014) and the evaluation of aversive and appetitive environments (Harris, McClure, van den Bos, Cohen, & Fiske, 2007; Mars et al. 2012; Winecoff et al. 2013; Veselic et al. 2023) in the latter. However, these relative specializations are embedded in a distributed network of areas that are recruited simultaneously (Braga and Leech 2015), warning against one-to-one matching of high-level functions with individual cortical areas (Hayden 2023).

Second, iFG function may be understood within a larger framework of numerous studies that document its modulation in social-emotional as well as cognitive paradigms. In the neuroimaging of social cognition, iFG has been noted to be active when violations of social expectations impose a reassessment of the interaction (Sanfey et al. 2003; Montague and Lohrenz 2007). King-Casas et al. (2008) demonstrated in a formalized strategy game that healthy controls activated the anterior insula/frontal operculum more than patients with borderline personality disorder (BPD) when attempting to restore the relationship by showing renewed cooperation efforts (see also Dziobek et al. 2011; Mier et al. 2013; Sosic-Vasic et al. 2019), suggesting a role in forming sophisticated representations of social interactions. This activity may be associated with the capacity of healthy individuals to form mental models of their interaction partners during the game, in contrast to BPD patients. Also in this region, however, it is possible to point out the existence of studies demonstrating a generic role in semantic disambiguation (Demb et al. 1995; Thompson-Schill et al. 1997; Binder and Desai 2011) that goes beyond social cognition.

In conclusion, our findings are consistent with those reported in meta-analyses of the cyberball paradigm, confirming its replicability. However, they are also consistent with those of a much broader set of studies. On the one hand, this draws attention to the relative lack of specificity of this paradigm and its findings, in line with the criticism of Cacioppo et al. (2013) and Mwilambwe-Tshilobo and Spreng (2021). The risk is one of treating operational constructs as if they were natural entities with a specific mapping onto cortical neurobiological processes. On the other hand, it underscores the internal consistency of neuroimaging data, when interpreted in an ecumenical approach, i.e. across the boundaries of traditional paradigm distinctions. As in other studies, the left iFG was recruited when violations of assumptions increased processing demands in interpreting the semantics of the social interaction.

## Methods

### Recruitment and image acquisition

The study was conducted at the Psychiatry and Psychotherapy Clinic of the University of Ulm, Germany, after approval by the Ethical Review Board. Healthy volunteers (N=27) were recruited from the local university. All participants were recruited through fliers distributed in the city of Ulm. Exclusion criteria were medical, neurological or psychiatric disorders. One participant did not complete the study, giving a final sample of N=26 participants (16 females, mean age 24.6, standard deviation 6.1).

Magnetic resonance imaging data were acquired using the ASL sequence described in Boland et al. (2018) using a 3-Tesla MAGNETOM Prisma scanner equipped with a standard 64-channel head/neck coil (Siemens, Erlangen, Germany) at the Department of Psychiatry of the University of Ulm. After positioning in the scanner, the heads of participants were padded to minimize movement artifacts during data acquisition. Participants could always communicate with the experimenter and had the option to interrupt the scanning session. Visual stimuli were presented on a 32-inch LCD screen (NordicNeuroLab AS, Bergen, Norway) positioned behind the scanner, viewed through a mirror attached to the head coil. The ASL sequence was applied with TR/TE: 4100/23.6 ms, matrix 64 × 64, field-of-view (FOV) 224 mm, pixel spacing 3.75 × 3.75 mm, slice thickness: 5 mm, 26 slices, flip angle 90°, PAT factor 2 (GRAPPA mode), bandwidth 2298 Hz/pixel, spin labelling phase 2400 ms, post-labeling delay 1000 ms. Conversion to CBF gave a volume every 8.2 seconds.

### Experimental task

The cyberball game, programmed using the Presentation® software package (Neurobehavioral Systems, Inc., Berkeley, CA), was designed to simulate social inclusion and exclusion through virtual ball-tossing between three cartoon player representations. The cyberball game task consisted of three conditions that were presented in a fixed order during the first run: (1/watch condition) passive viewing, where participants watched the ball toss without interacting, (2/inclusion play condition) social inclusion, where participants actively participated by throwing the ball to one of the virtual players, and (3/exclusion condition) social exclusion, where participants initially played as in the inclusion condition but were subsequently excluded. Each scan started with an image displaying two virtual players in the upper corners of the screen and an arm symbolizing the participant located at the bottom center.

After an initial rest period of 3 sec, the first throw occurred between 500 and 1000 milliseconds after the start of the game to increase the realism of the social interaction. Each trial type, such as a left-to-right throw, consisted of eight 200 ms stimulus events, totalling 1600 ms per trial for each condition. Both the watch and inclusion conditions were limited to a duration of 2 minutes. The exclusion condition was extended to 2 min 30 sec to ensure that only experienced exclusion could be separated from inclusion in subsequent analyses. The exclusion phase began immediately after a 20 sec. inclusion phase and lasted until the end of the block. Throughout the experiment, participants responded using a button box, allowing them to choose between playing as the right or left player.

### Statistical modelling and analysis

Images were realigned prior to computing estimates of cerebral blood flow with equation [1] in Wang et al. (2003). Mean realigned EPI images were used to compute estimates of registration to a MNI template, which were subsequently applied to the CBF images. Finally, the registered CBF images (resampling size: 2mm isotropic) were smoothed with a Gaussian kernel (FWHM 8mm).

At the first level, conditions were modelled as blocks, all comprising 14 CBF volumes (the first three volumes of exclusion block, which lasted 24.6sec longer than the others, were modelled as a confounder, so as to model exclusion with a block of the same length as the other blocks and only considering when it could become clear that the participants was being excluded). To adjust for physiological noise, the mean activity and the first 7 principal components from white matter and ventricles (8 components in total, Behzadi et al. 2007), and the mean activity from cranial bone (Huber et al. 2024) were added as covariates to the model. The segments were extracted from the segmentation computed by SPM as part of the registration algorithm. To avoid partial volume effects, activity was extracted from registered volumes (i.e. voxel size 2mm) without smoothing. Cranial bone was eroded by 1 voxel (to avoid sampling subdural space) and white matter by 2 voxels. Ventricles were selected from the CSF segment by masking it with a priori maps of ventricles from the Harvard-Oxford Cortical and Subcortical Atlas (https://fsl.fmrib.ox.ac.uk/fsl/fslwiki/Atlases). Contrasts of interest (play vs. watch, exclusion vs. play, exclusion vs. watch) were brought to the second level to account for subjects as a random factor. Anatomical regions of interest for the iFG and vACC were defined with the aal atlas (iFG: 1256 voxels; vACC: 1498 voxels).

## Acknowledgments

We would like to thank Eun-Jin Sim, of the Psychiatry Clinic of the University of Ulm, for coding the cyberball game for use in the scanner and making it available to us. Data collection was supported by a Neuron-ERANET grant (project ‘BrainCYP’, grant number BMBF 01EW1402B) to RV.

## Appendix

**Table A1.**
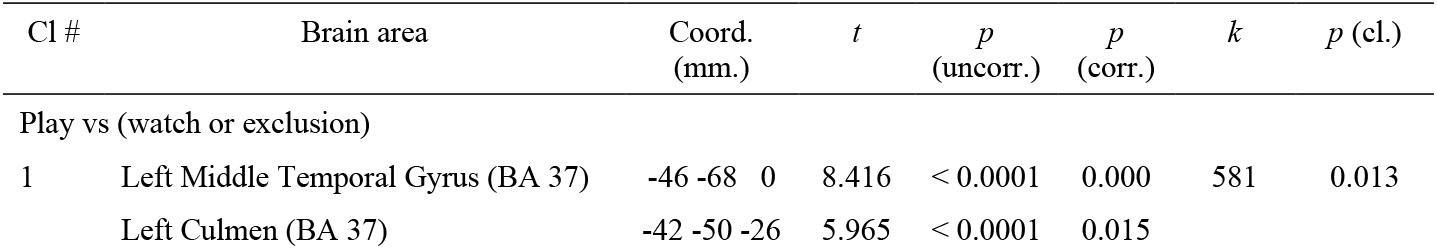
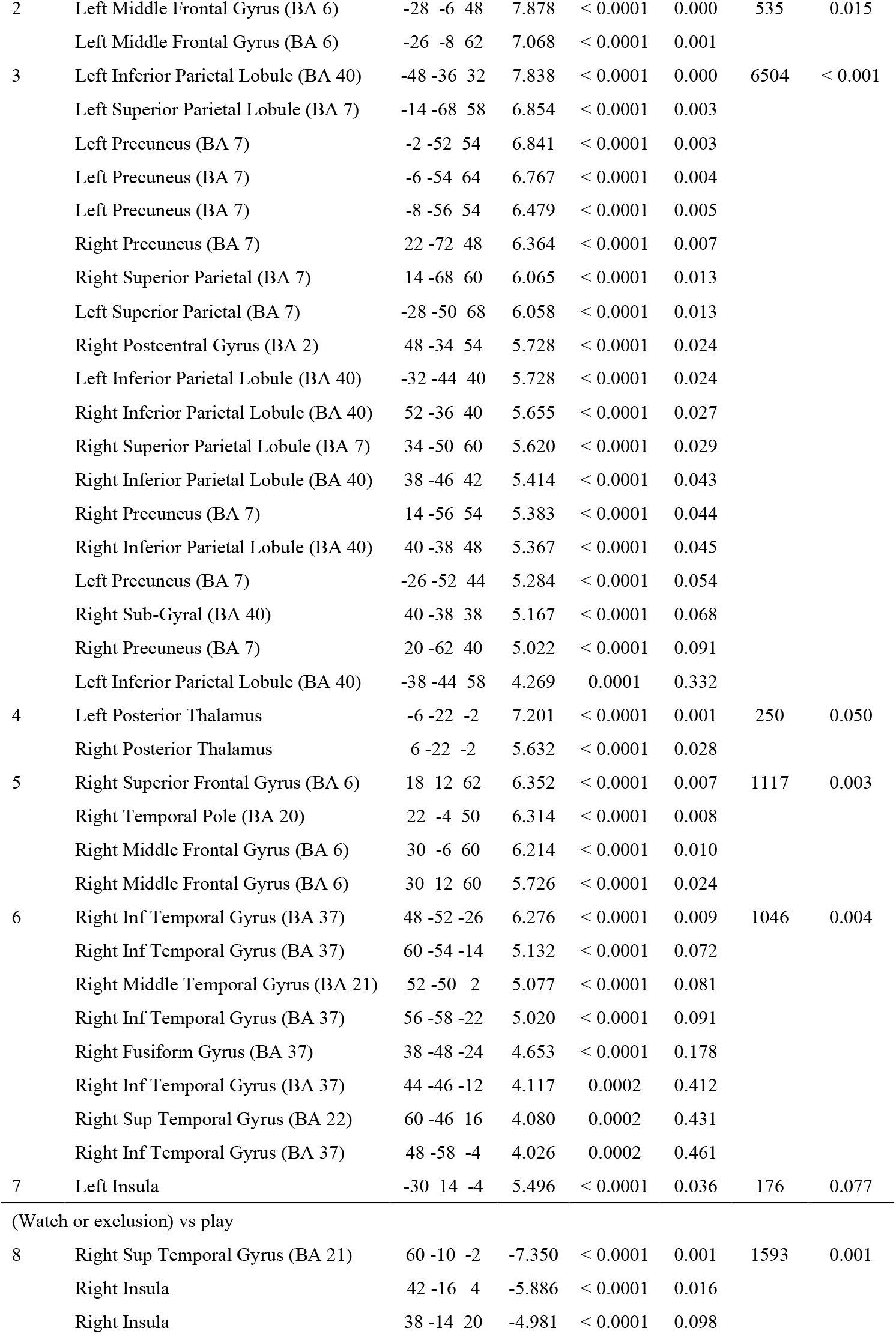
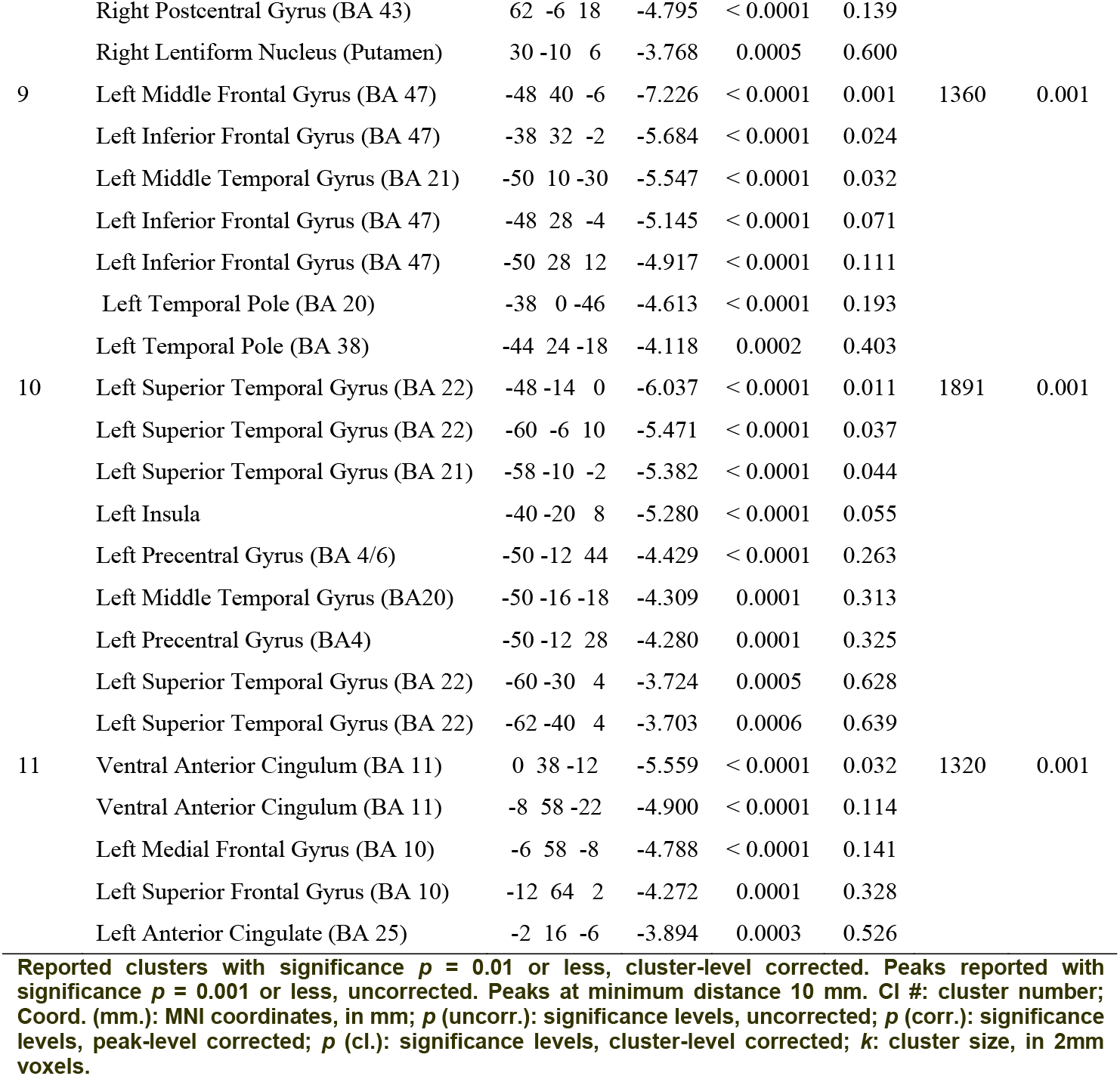
Contrast play vs (watch or exclusion)

**Table A2.**
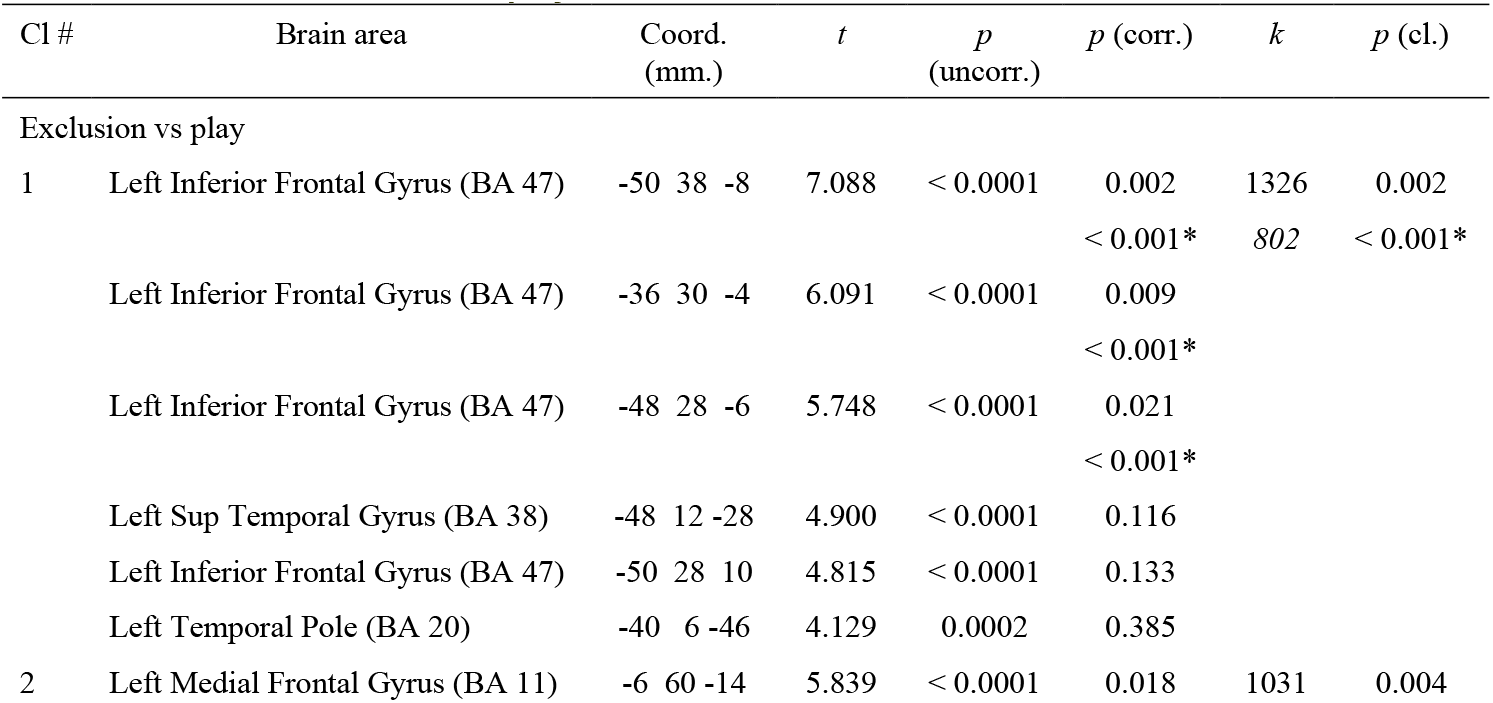
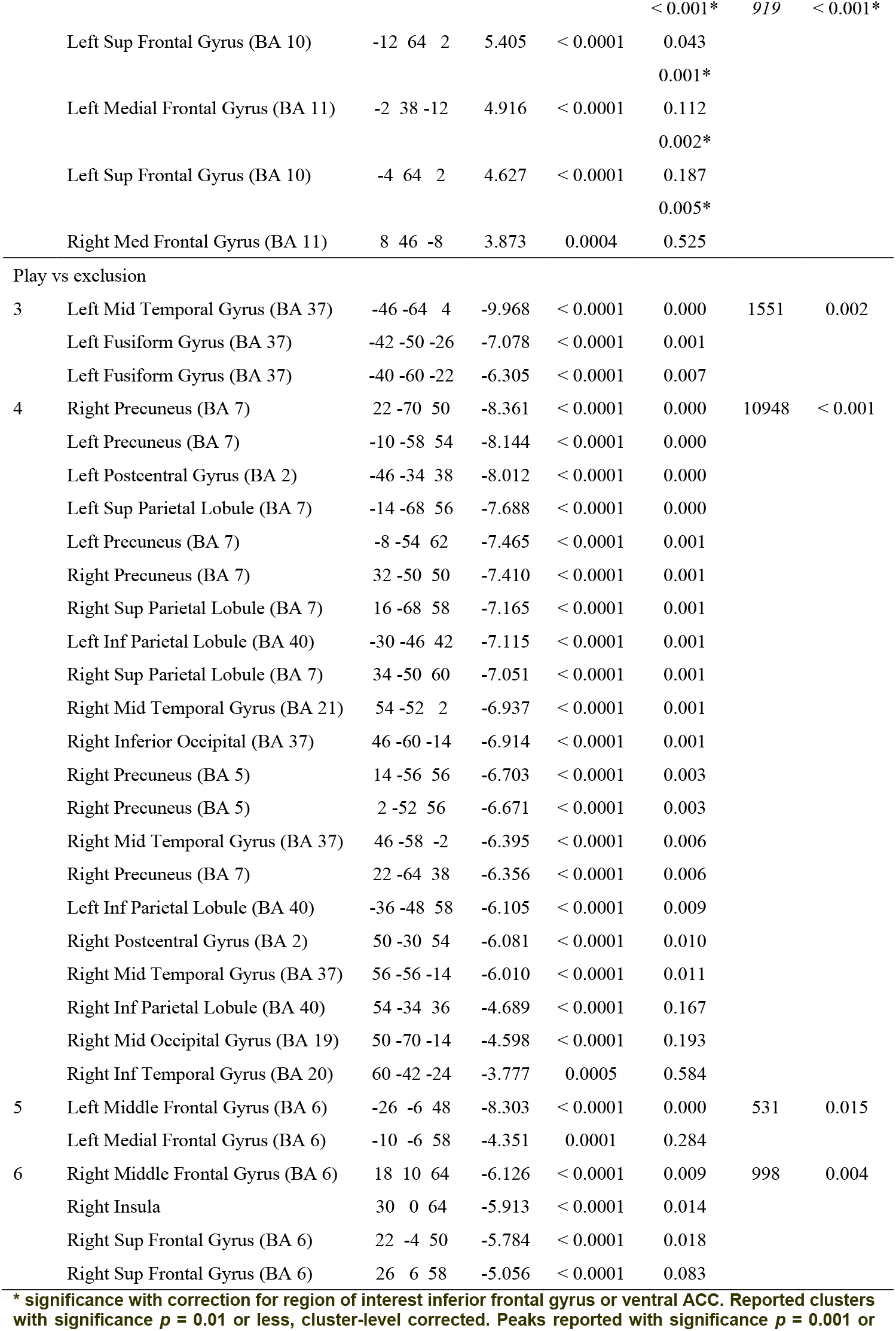
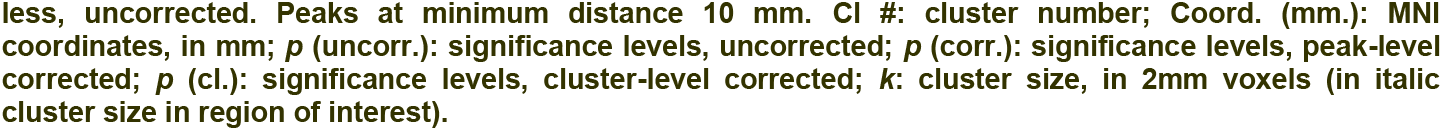
Contrast exclusion vs play.

**Table A3.**
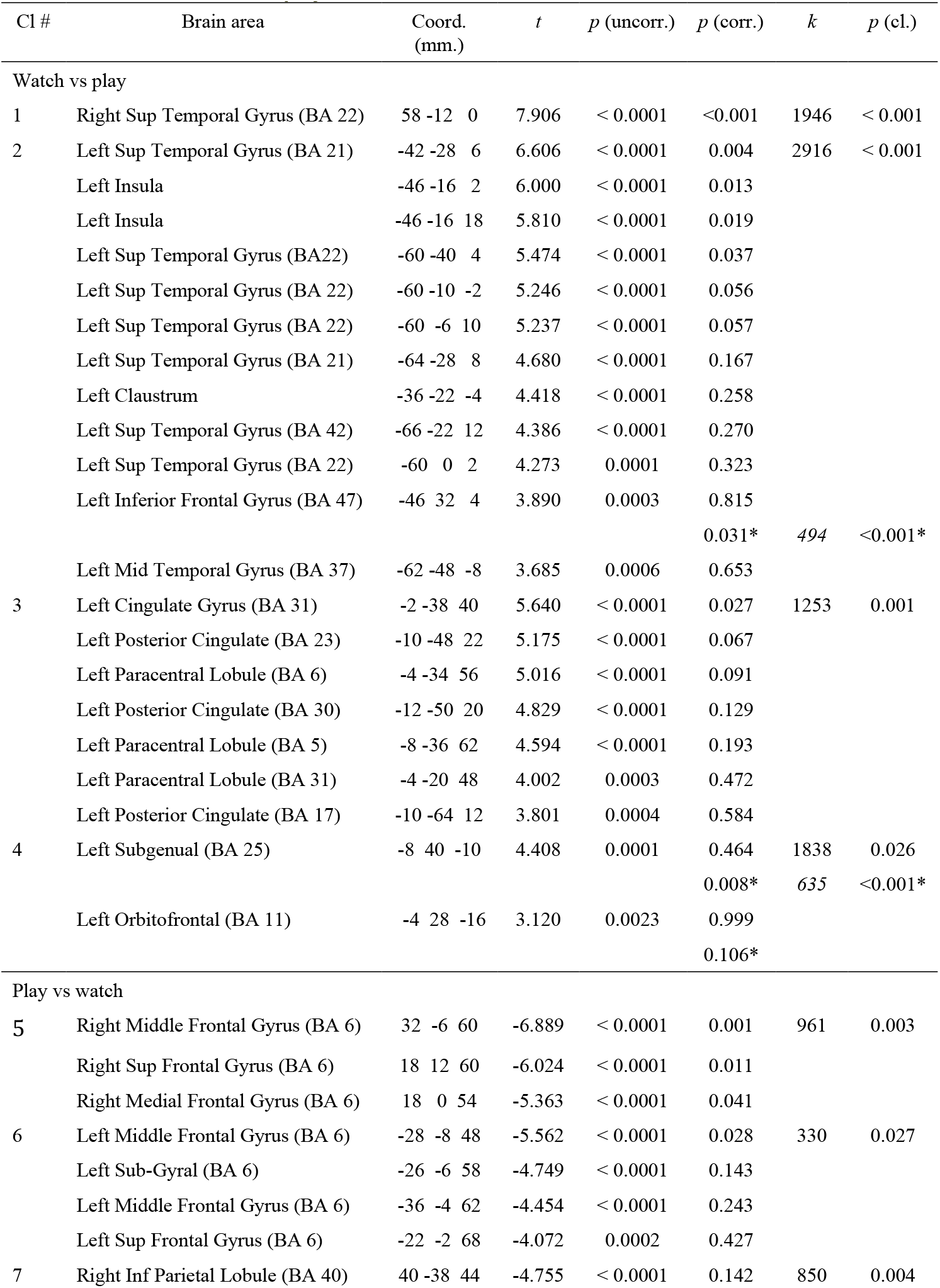
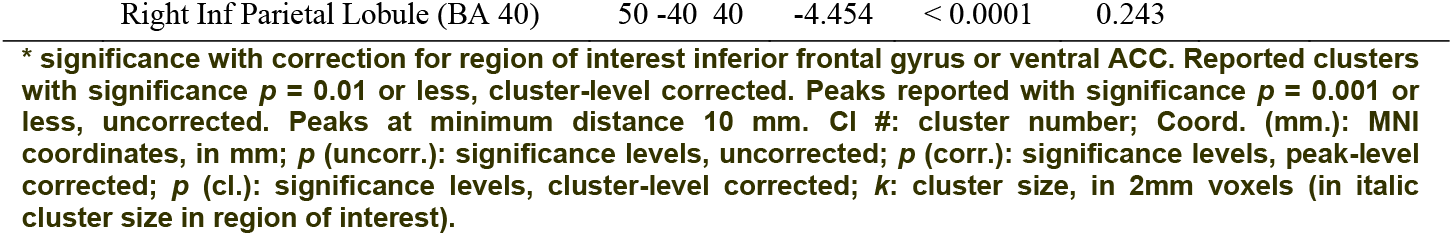
Contrast watch vs play.

**Table A4.** Contrast exclusion vs watch.

| CI # | Brain area | Coord.<br>(mm.) | $t$ | $p$<br>(uncorr.) | $p$ (corr.) | $k$ | $p$ (cl.) |
| --- | --- | --- | --- | --- | --- | --- | --- |
| Exclusion vs watch |  |  |  |  |  |  |  |
| 1 | Left Superior Frontal Gyrus (BA 11) | -16 58 -18 | 3.936 | < 0.001 | 0.489 | 18 | 0.430 |
| 2 | Left Inferior Frontal Gyrus (BA 47) | -34 28 -6 | 3.582 | < 0.001 | 0.699 | 6 | 0.599 |
|  |  |  |  |  | 0.031* | 50 | 0.053* |
| 3 | Left Ventromedial Prefrontal (BA 11) | -10 60 -14 | 3.740 | < 0.001 | 0.027* | 67 | 0.050* |
| Watch vs exclusion |  |  |  |  |  |  |  |
| 4 | Left Superior Temporal Gyrus(BA 42) | -64 -34 18 | -7.602 | < 0.001 | 0.001 | 3615 | <0.001 |
| 5 | Right Superior Parietal Lobule (BA5) | 18 -58 62 | -7.306 | < 0.001 | 0.001 | 2949 | <0.001 |
| 6 | Right Insula(BA 13) | 40 -20 6 | -5.307 | < 0.001 | 0.047 | 1220 | 0.003 |
| 7 | Right Fusiform (BA 19) | 40 -72 -20 | -4.928 | < 0.001 | 0.099 | 1249 | 0.003 |
| 8 | Left Superior Frontal (BA 6) | -24 -8 54 | -4.466 | < 0.001 | 0.228 | 36 | 0.309 |
| 9 | Left Middle Occipital Gyrus (BA 39) | -42 -76 20 | -4.238 | < 0.001 | 0.331 | 44 | 0.275 |

\* significance with correction for region of interest inferior frontal gyrus or ventral ACC. Peaks reported with significance $p = 0.001$ or less, uncorrected. Peaks at minimum distance 10 mm. CI #: cluster number; Coord. (mm.): MNI coordinates, in mm; $p$ (uncorr.): significance levels, uncorrected; $p$ (corr.): significance levels, peak-level corrected; $p$ (cl.): significance levels, cluster-level corrected; $k$ : cluster size, in 2mm voxels (in *italic cluster size in region of interest*).

## References

Arana, F.S., Parkinson, J.A., Hinton, E., Holland, A.J., Owen, A.M., Roberts, A.C., 2003. Dissociable contributions of the human amygdala and orbitofrontal cortex to incentive motivation and goal selection. J. Neurosci. 23, 9632–9638.

Bassett, D.S., Bullmore, E., 2006. Small-world brain networks. Neurosicenstist 12, 512–523.

Behzadi, Y., Restom, K., Liau, J., Liu, T.T., 2007. A component based noise correction method (CompCor) for BOLD and perfusion based fMRI. NeuroImage 37, 90–101.

Binder, J.R., Desai, R.H., 2011. The neurobiology of semantic memory. Trends Cogn. Sci. 15, 527–536.

Binder, J.R., Desai, R.H., Graves, W.W., Conant, L.L., 2009. Where is the semantic system? A critical review and meta-analysis of 120 functional neuroimaging studies. Cereb. Cortex 19, 2767–2796.

Boland, M., Stirnberg, R., Pracht, E., Kramme, J., Viviani, R., Stingl, J.C., Stöcker, T., 2018. Accelerated 3D-GRASE imaging improves quantitative multiple post-labeling delay arterial spin labeling. Magn. Reson. Med. 80, 2475–2484.

Braga, R.M., Leech, R., 2015. Echoes of the brain: Long-scale representation of whole-brain functional networks within transmodal cortex. Neurosicenstist 21, 540–551.

Cacioppo, S., Frum, C., Asp, E., Weiss, R.M., Lewis, J.W., Cacioppo, J.T., 2013. A quantitative meta-analysis of functional imaging studies of social rejection. Sci. Reports 3, 2027.

Corbetta, M., Patel, G., Shulman, G.L., 2008. The reorienting system of the human brain: From environment to theory of mind. Neuron 58, 306–324.

Davis, K.D., Taylor, S.J., Crawley, A.P., Wood, M.L., Mikulis, D.J., 1997. Functional MRI of pain- and attention-related activations in the human cingulate cortex. J. Neurophysiol. 77, 3370–3380.

Decety, J., 2010. To what extent is the experience of empathy mediated by shared neural circuits? Emotion Rev. 2, 204–207.

Demb, J.B., Desmond, J.E., Wagner, A.D., Vaidya, C.J., Glover, G.H., Gabrieli, J.D.E., 1995. Semantic encoding and retrieval in the left inferior prefrontal cortex: A functional MRI study of task difficulty and process specificity. J. Neurosci. 15, 5870–5878.

Dziobek, I., Preißler, S., Grozdanovic, Z., Heuser, I., Heekeren, H.R., Roepke, S., 2011. Neuronal correlates of altered empathy and social cognition in borderline personality disorder. NeuroImage 57, 539–548.

Eisenberger, N.I., Liebermann, M.D., Williams, K.D., 2003. Does rejection hurt? An fMRI study of social exclusion. Science 203, 290–292.

FitzGerald, T.H.B., Friston, K.J., Donal, R.J., 2012. Action-specific value signals in reward-related regions of the human brain. J. Neurosci. 32, 16417–16423.

Harris, L.T., McClure, S.M., van den Bos, W., Cohen, J.D., Fiske, S.T., 2007. Regions of the MPFC differntially tuned to social and nonsocial affective evaluation. Cogn. Aff. Behav. Neurosci. 7, 309–316.

Hayden, B.Y., 2023. The dangers of cortical brain maps. J. Cogn. Neurosci. 35, 372–375.

Huber, D., Rabl, L., Orsini, C., Labek, K., Viviani, R., 2024. The fMRI global signal and its association with the signal from cranial bone. NeuroImage 297, 120754.

Kiefer, M., Sim, E.J., Heil, S., Brown, R., Herrnberger, B., Spitzer, M., Grön, G., 2021. Neural signatures of bullying experience and social rejection in teenagers. PLoS One 10.1371/journal.pone.0255681.

King-Casas, B., Sharp, C., Lomax-Bream, L., Lohrenz, T., Fonagy, P., Montague, P.R., 2008. The rupture and repair of cooperation in borderline personality disorder. Science 321, 806–810.

Kross, E., Berman, M.G., Mischel, W., Smith, E.E., Wager, T.D., 2011. Social rejection shares somatosensory representations with physical pain. Proc. Natl Acad. Sci. USA 108, 6270–6275.

Labek, K., Berger, S., Buchheim, A., Bosch, J., Spohrs, J., Dommes, L., Beschoner, P., Stingl, J.C., Viviani, R., 2017. The iconography of mourning and its neural correlates: A functional neuroimaging study. Soc. Cogn. Aff. Neurosci. 12, 1303–1313.

Labek, K., Sittenberger, E., Kienhöfer, V., Rabl, L., Messina, L., Schurz, M., Stingl, J.C., Viviani, R., 2023. The gradient model of brain organization in decisions involving ‘empathy for pain’. Cereb. Cortex 33, 5839–5850.

Margulies, D.S., Ghosh, S.S., Goulas, A., Falkiewicz, M., Huntenburg, J.M., Langs, G., Bezgin, G., Eickhoff, S.B., Castellanos, F.X., Petrides, M., Jefferies, E., Smallwood, J., 2016. Situating the default-mode network along a principal gradient of macroscale cortical organization. Proc. Natl Acad. Sci. USA 113, 12574–12579.

Mars, R.B., Neubert, F.X., Noonan, M.A.P., Sallet, J., Toni, I., Rushworth, M.F.S., 2012. On the relationship between the ‘default mode network’ and the ‘social brain’. Front. Hum. Neurosci. doi: 10.3389/fnhum.2012.00189.

Masten, C.L., Eisenberger, N.I., Borofsky, L.A., McNealy, K., Pfeifer, J.H., Dapretto, M., 2011. Subgenual anterior cingulate responses to peer rejection: A marker of adolescents’ risk for depression. Dev. Psychopathol. 23, 283–292.

Messina, I., Sambin, M., Beschoner, P., Viviani, R., 2016. Changing views of emotion regulation and neurobiological models of the mechanism of action of psychotherapy. Cogn. Aff. Behav. Neurosci. 16, 571–587.

Mesulam, M.-M., 1990. Large-scale neurocognitive networks and distributed processing for attention, language, and memory. Ann. Neurol. 28, 597–613.

Mier, D., Lis, S., Esslinger, C., Sauer, C., Hagenhoff, M., Ulferts, J., Gallhofer, B., Kirsch, P., 2013. Neuronal correlates of social cognition in borderline personality disorder. Soc. Cogn. Aff. Neurosci. 8, 531–537.

Montague, P.R., Lohrenz, T., 2007. To detect and correct: Norm violations and their enforcement. Neuron 56, 14–18.

Mwilambwe-Tshilobo, L., Spreng, N., 2021. Social exclusion reliably engages the dafault network: A meta-analysis of Cyberball. NeuroImage 227, 117666.

Panksepp, J., 2003. Feeling the pain of social loss. Science 302, 237–239.

Rotge, J.Y., Lemogne, C., Hinfray, S., Huguet, P., Grynszpan O Tartour, E., George, N., Fossati, P., 2014. A meta-analysis of the anterior cingulate contribution to social pain. Soc. Cogn. Aff. Neurosci. 10, 19–27.

Rudebeck, P.H., Murray, E.A., 2014. The orbitofrontal oracle: Cortical mechanisms for the prediction and evaluation of specific behavioral outcomes. Neuron 84, 1143–1156.

Sanfey, A.G., Rilling, J.K., Aronson, J.A., Nystrom, L.E., Cohen, J.D., 2003. The neural basis of economic decision-making in the ultimatum game. Science 300, 1755–1758.

Sawamoto, N., Honda, M., Okada, T., Hanawaka, T., Kanda, M., Fukuyama, H., Konisci, J., Shibasaki, H., 2000. Expectation of pain enhances responses to nonpainful somatosensory stimulation in the anterior cingular cortex and parietal operculum/posterior insula: An event-related functional magnetic resonance imaging study. J. Neurosci. 20, 6438–7445.

Schurz, M., Radua, J., Tholen, M.G., Maliske, L., Margulies, D.S., Mars, R.B., Sallet, J., Kanske, P., 2021. Toward a hierarchical model of social cognition: A meta-analysis and integrative review of empathy and theory of mind. Psychol. Bull. 147, 293–327.

Somerville, L.H., Heatherton, T.F., Kelley, W.M., 2006. Anterior cingulate cortex responds differentially to expectancy violation and social rejection. Nat. Neurosci. 9, 1007–1008.

Sosic-Vasic, Z., Eberhard, J., Bosch, J.E., Dommes, L., Labek, K., Buchheim, A., Viviani, R., 2019. Mirror neuron activations in encoding of psychic pain in broderline personality disorder. NeuroImage Clin. 22, 101737.

Sporns, O., Honey, C.J., Kötter, R., 2007. Identification and classification of hubs in brain networks. PLoS One 2, e1049.

Thompson-Schill, S.L., D’Esposito, M., Aguirre, G.K., Farah, M.J., 1997. Role of left inferior prefrontal cortex in retrieval of semantic knowledge: A re-evaluation. Proc. Natl Acad. Sci. USA 94, 14792–14797.

van den Heuvel, M.P., Sporns, O., 2013. Network hubs in the human brain. Trends Cogn. Sci. 17, 683–696.

Veselic, S., Muller, T.H., Gutierrez, E., Behrens, T.E., Hunt, L.T., Butler, J.L., Kennerley, S.W., 2023. A cognitive map for value-guided choice in ventromedial prefrontal cortex. bioRxiv doi: 10.1101/2023.12.15.571895.

Vijayakumar, N., Cheng, T.W., Pfeifer, J.H., 2017. Neural correlates of social exclusion across ages: A coordinate-based meta-analysis of functional MRI studies. NeuroImage 153, 359–368.

Viviani, R., Dommes, L., Bosch, J.E., Labek, K., 2020. Segregation, connectivity, and gradients of deactivation in neural correlates of evidence in social decision making. NeuroImage 223, 117339.

Vogt, B.A., Finch, D.M., Olson, C.R., 1992. Functional heterogeneity in cingulate cortex. The anterior executive and posterior evaluative regions. Cereb. Cortex 2, 435–443.

Wang, J.J., Alsop, D.C., Li, L., Listerud, J., Gonzalez-At, J.B., Detre, J.A., 2003. Arterial transit time imaging with flow encoding arterial spin tagging (FEAST). Magn. Reson. Med. 50, 599–607.

Williams, K.D., Cheung, C.K.T., Choi, W., 2000. Cyberostracism: Effects of being ignored over the internet. J. Pers. Soc. Psychol. 79, 748–762.

Winecoff, A., Clithero, J.A., McKell Carter, R., Bergman, S.R., Wang, L., Huettel, S.A., 2013. Ventromedial prefrontal cortex encodes emotional value. J. Neurosci. 33, 11032–11029.

